# Beyond Bonferroni Revisited: Concerns over inflated false positives in conservation genetics, genetics, and neuroscience

**DOI:** 10.1101/408997

**Authors:** Tonya White, Jan van der Ende, Thomas E. Nichols

**Affiliations:** Department of Child and Adolescent Psychiatry, Erasmus University Medical Center, Rotterdam, the Netherlands; Department of Radiology, Erasmus University Medical Center, Rotterdam, the Netherlands; Oxford Big Data Institute, Li Ka Shing Centre for Health Information and Discovery, Nuffield Department of Population Health, University of Oxford, Oxford, OX3 7LF, UK; Wellcome Centre for Integrative Neuroimaging, FMRIB, Nuffield Department of Clinical Neurosciences, University of Oxford, Oxford, OX3 9DU; Department of Statistics, University of Warwick, Coventry, CV4 7AL, UK

**Author notes:** **Corresponding Author** Tonya White, M.D., Ph.D., M.Sc. Eng, Associate Professor, Department of Child and Adolescent Psychiatry, Erasmus MC-Sophia / Kamer KP-2869, Postbus 2060, 3000 CB Rotterdam.

**Keywords:** Multiple Testing Correction, False Discovery Rate, Family Wise Error, Benjamini Hochberg, Benjamini Yekutieli

## Abstract

In 2006, Narum published a paper in Conservation Genetics that was motivated by the stringent nature of the Bonferroni approach for family wise error correction. That work suggested that the approach of Benjamini and Yekutieli in 2001 provided adequate correction and was more biologically relevant. However, there are crucial differences between the original Benjamini and Yekutieli procedure and that described by Narum. After carefully reviewing both papers, we believe that the Narum procedure is both different than the Benjamini and Yekutieli procedure and does not adequately control for family wise error. We provide an overview of approaches for FWE correction as well as evidence for the faulty implementation of the Benjamini and Yekutieli procedure by Narum using the equations from the respective papers, data from both papers, and the results of simulation.

## Introduction

In 2006, Narum published a paper in Conservation Genetics motivated by the stringent nature of the Bonferroni approach for multiple testing correction, suggesting the False Discovery Rate (FDR) method proposed by (Benjamini and Yekutieli 2001) as an alternative that is both powerful but also more biologically relevant. His paper titled “Beyond Bonferroni: Less conservative analyses for Conservation Genetics” has been cited over 500 times [https://link.springer.com/article/10.1007/s10592-005-9056-y]. The article has not only been cited in the field of conservation genetics, but also has been increasingly cited in the fields of medicine and neuroscience. These studies apply the approach of Narum (2006) attributed to the Benjamini and Yekutieli (2001) (BY) procedure for muliple testing correction.

However, a careful review of the published BY approach and what Narum describes as the BY method, there are crucial differences. Due to an omission of one term, Narum’s implementation of BY is incorrect and cannot be guaranteed to control the FDR. Thus, we believe that the Narum publication has created confusion about the BY procedure and its misuse is being propogated along an increasing number of studies. Thus, we have two goals of this paper: The first is to provide an overview of the Bonferroni method, the original Benjamini & Hochberg (2000) FDR (BH-FDR), and BY’s method (BY-FDR); the second goal is to describe faulty implimentation of the BY-FDR approach described by Narum. We will demonstrate that using the multiple testing correction described by Narum results in an excessive number of false positives, especially when a larger number of multiple tests are performed.

## Theory

We first review the different multiple testing approaches discussed by Narum (2006) using his notation as closely as possible. For a collection of *k* tests, each with a corresponding p-value, *p*_*i*_, *i*=1,…,*k*. A multiple testing procedure identifies a subset of the *k* tests as significant while controlling for some measure of false positive risk that takes into account the number of tests performed. The Bonferroni method controls the family-wise error (FWE), the chance of one or more false positives, by using a fixed threshold of:

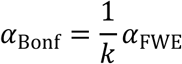

where *α*_FWE_ is the desired FWE level: All tests with *p* _*i*_ ≤ *α*_Bonf_ can be declared significant while controlling the FWE.

Benjamini & Hochberg (2000) introduced the False Discovery Rate (FDR) for multiple testing correction. In describing the FDR it is useful to first define the false discovery proportion (FDP): FDP is the ratio of the number of false positive tests to total number of significant tests, defined as 0 if no tests are significant. The FDR is the expected value of FDP; put another way, FDR is the expected proportion of false positives among positives. To find FDR-significant tests, denote the ordered p-values *p* _(1)_ ≤ *p* _(2)_) ≤…≤ *p* _(*k*)_. Then for a desired *α*_FDR_, let the index *i* be found as

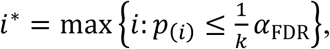

and the tests with *p* _*i*_ ≤ *p*(_*i\**_) can be declared significant while controlling FDR at *α*_FDR_.

The assumptions of this Benjamini & Hochberg FDR procedure (BH-FDR) are independence among the test statistics (Benjamini & Hochberg, 2000). However, BY found that weaker assumptions could be used, allowing a general form of positive dependence among the test statistics. The BY work, however, also proposed another method for controlling FDR that makes *no assumptions* about the dependence among the tests, as long as a more stringent criterion was used (Theorem 1.3, BY), with the index ^*i* *^_BY_ computed:

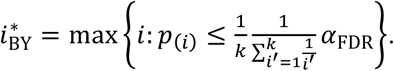

With this approach, the tests with 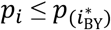 are marked significant and FDR is controlled at *α* _FDR_ under any form of dependency. Notably 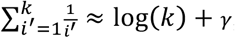 where λ ≈ 0.57721 is Euler’s constant. This is the method we refer to by BY-FDR.

We can now make a quick comparison of three methods on the basis of the smallest p-value *p* _(1)_: Bonferroni has the fixed threshold *α*_FWE_/*k*, while BH-FDR will compare *p*_(1)_ to *α*_FDR_/*k* and BY-FDR will compare *p*_(1)_ to approximately *α*_FDR_/(*k* log(*k*)). Of course, BH-FDR and BY-FDR are adaptive and compare increasing p-values to successively more lenient thresholds, but this comparison for *p*_(1)_ points to how BY-FDR is much more stringent than BH-FDR.

Now, in Narum (2006), the author incorrectly states that the BY-FDR threshold is fixed and equal to:

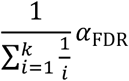

This is a fundamental error, as a key feature of FDR methods is that they are adaptive. The error arose from neglecting that the equation above was merely one component of the BY procedure (to be substituted for q in B-Y Eq. (1) on pp. 1167 (Benjamini and Yekutieli 2001)). The Narum procedure results in a fixed threshold for a specific *k*.

Since a fixed threshold specifies the average or per comparison error rate (PCE), we can assess the impact of this error. Assuming the complete null, i.e. no signal for any test, *k* × PCE is the expected number of false positives. For the threshold at the 0.05 level, for *k* = 105, *k* × PCE ≈ 1, while for *k* = 1590, *k* × PCE ≈ 10. This demonstrates that Narum’s result can be assured to produce an increasing number of false positives for an increasing *k*. In contrast, for Bonferroni *k* × PCE is exactly *α*_FWE_, i.e. always less than 1, and every valid FWE or FDR level α procedure is guaranteed to produce *no* false positives with probability 1-α (again, in this complete null setting). While the Narum approach does asymptote to zero as *k* approaches infinity, it approaches zero extremely slowly. For example, with 10 million tests performed, the Narum p-value threshold is 0.003, in contrast to the Bonferroni threshold of 0.000000005.

To evaluate the rate of significant p-values between the Bonferroni, B-H, B-Y, and Narum’s interpretation of the B-Y approach we conducted a simulation. We created 50,000 random realizations where random p-values were computed from a standard Normal distribution. We considered *k* ranging from 1 to 30 tests, where all tests were independent, and used nominal *α*_FWE_ = α_FDR_ = 0.05 for all methods. In this null setting, any “discovery” is a false discovery and so measured FDR and FWE will be the same. So we computed the proportion of realizations where any p-values we found significant (FDR & FWE), as well as the proportion of tests among the k that were false positives, known as the Per Comparison Error rate (PCE). Python code is available in Appendix 1.

Figure 1 shows the FDR and FWE as a function of the number of tests, showing that Bonferroni and BH-FDR both control false positives as expected (as an aside, while Bonferroni is often regarded as conservative, in this setting of small *k* and independent tests, it is essentially exact). The FDR/FWE of BY-FDR becomes increasing conservative while Narum’s method has inflated false positives even for k=2 tests, and has a near linear increase with increasing *k*. In all realizations there was never more than 1 detection, and hence the PCE was identical to the FDR/FWE (not shown).

**Figure 1.**
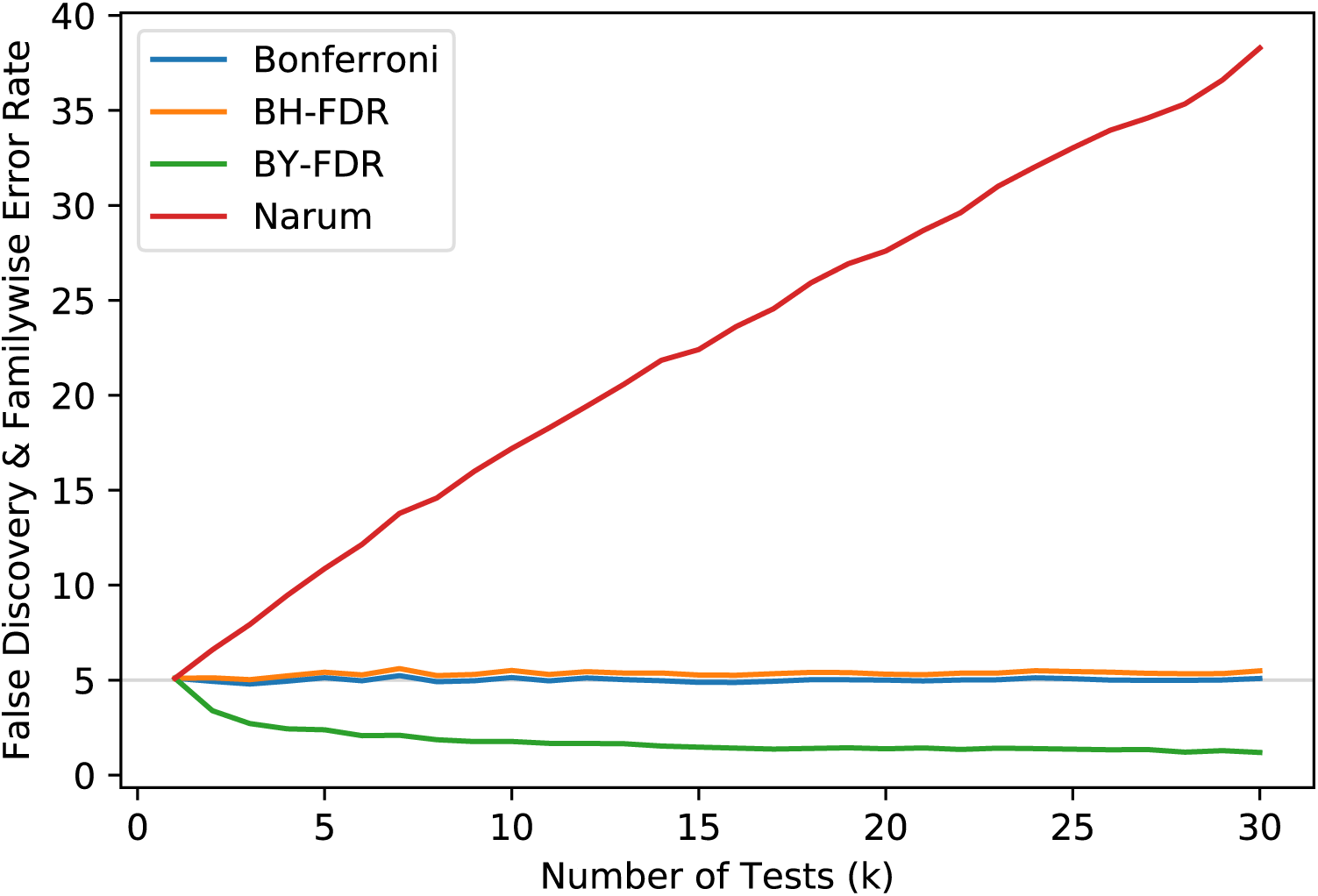
Percentage of false positive results compared to the number of independent tests performed. The results were derived from computer simulations comparing four different approaches for multiple testing correction. Simulations were performed in python and utilized a 5000 iterations of a random Normal distribution converted to p values.

We also consider the specific set of 15 p-values used in Narum (2006), tabulating the p-value threshold that were used for significance testing for each of the four methods. Table 1 shows the thresholds used for each of the 15-exemplar p-values, with significant tests marked in bold. It can be seen that the BY-FDR and the Narum approach are not the same, with Narum finding 4 significant tests as compared to BY-FDR’s having two significant tests.

**Table 1.** A set of p-values from 15 significance testing taken from the Narum 2006 paper and comparison with four approaches to multiple testing. Numbers in **bold** reflect those in the p-value row that are significant.

| p-value<br>Examples | Bonferroni | Benjamini &<br>Hochberg | Benjamini &<br>Yekutieli | Narum |
| --- | --- | --- | --- | --- |
| 0.0001 | **0.0033** | **0.0033** | **0.0010** | **0.0151** |
| 0.0010 | **0.0033** | **0.0067** | **0.0020** | **0.0151** |
| 0.0062 | 0.0033 | **0.0100** | 0.0030 | **0.0151** |
| 0.0101 | 0.0033 | **0.0133** | 0.0040 | **0.0151** |
| 0.0214 | 0.0033 | **0.0167** | 0.0050 | 0.0151 |
| 0.0227 | 0.0033 | **0.0200** | 0.0060 | 0.0151 |
| 0.0273 | 0.0033 | **0.2333** | 0.0070 | 0.0151 |
| 0.0292 | 0.0033 | **0.0267** | 0.0080 | 0.0151 |
| 0.0311 | 0.0033 | **0.0300** | 0.0090 | 0.0151 |
| 0.0323 | 0.0033 | **0.0333** | 0.0100 | 0.0151 |
| 0.0441 | 0.0033 | 0.0367 | 0.0111 | 0.0151 |
| 0.0490 | 0.0033 | 0.0400 | 0.0121 | 0.0151 |
| 0.0573 | 0.0033 | 0.0433 | 0.0131 | 0.0151 |
| 0.1262 | 0.0033 | 0.0467 | 0.0141 | 0.0151 |
| 0.5794 | 0.0033 | 0.0500 | 0.0151 | 0.0151 |

## Discussion

Approaches for multiple testing correction have been present for over half a century. In the late 1950’s, Olive Jean Dunn adapted the Italian mathematician Carlo Emilio Bonferroni’s theory of inequalities for use in statistics (Dunn 1961). However, since the Bonferroni approach is one of the most conservative, especially when the multiple tests are dependent, multiple alternative approaches to correct for multiple testing have surfaced. Two commonly used approaches include the Benjamini & Hochberg that was introduced in 1995 and the Benjamini and Yekutieli approach that was introduced in 2001. In 2006, Narum published a paper which provided an overview and examples of the BY-FDR procedure. However, we demonstrate that the approach described by Narum is an incorrect depiction of the BY-FDR approach.

We believe that Narum used an equation from the BY paper (shown above) out of context. A careful reading of Benjamini and Yekutieli (2001) finds that this expression (from Theorem 1.3 on pp. 1169 of BY) is to be entered as the *α* in the B-H equation (Eq. (1) on pp. 1167 in BY), producing an adaptive threshold. Further, we also show, based on a series of p-values taken also from the Narum paper (Table 1) that different results are obtained from the BY-FDR approach and the Narum approach.

Direct calculation shows that Narum’s incorrect implementation of BY-FDR has expected number of false positives that increases linearly with number of tests *k*, and invalid and increasing false positive rates that, crucially, are drastically different from BY-FDR’s valid but conservative performance. We believe that over 500 publications citing Narum (2006) are liable to have this inflated rate of false positives in their results. For example, we are particularly concerned about papers that cite both the BY paper as the method used, but use the multiple testing correction described by Narum 2006 (Chye et al.; Whittle et al. 2016; Woolley et al. 2018).

We do agree with Narum that the Bonferroni approach is often conservative for multiple testing correction, especially with dependent data. However, there has also been a growing concern that many studies fail to replicate (Ioannidis 2005; Open Science Collaboration 2015; Nichols et al. 2017). In the past, analyses were performed without adequately controlling for the numbers of tests performed (Carp 2012) which resulted in numerous type I errors. We know of no justification to use the procedure described by Narum for multiple testing, and are unaware of any formal metric of false positives that it controls. Thus, we would recommend that this approach not be used for multiple testing correction and the work corrected to properly impliment the BY-FDR approach.

## Supplemental Material

### Python Code for Simulation of the Multiple Testing

““”

Simulations performed on multiple testing in Python

Author: Tonya White, MD, PhD

Adapted by Tom Nichols, PhD on 3 September 2018

Date: 24 April 2018

Location: Rotterdam, Netherlands

This program performs simulations of multiple testing to test the Narum paper

““”

import numpy as np

import scipy.stats as st

import matplotlib as ml

import matplotlib.pyplot as plt

from matplotlib import cm

##################################################################################################

\# You can change variables within this section

m = 30

\# Maximum number of multiple tests to perform

k = 5000

\# Number of iterations per test

alpha = 0.05

####################################################################################

\# Create an array to hold the mean for all the simulations

bonf = np.zeros(m)

bh = np.zeros(m)

by = np.zeros(m)

narum = np.zeros(m)

\# Loop structure to perform iterative tests for percent of false positives

for x in range(m):

pvals = np.zeros(x+1)

bonthr = alpha / (1 + x)

\# Calculate the critical p value for the Narum value

ui = 0

for t in range(x+1):

ui = ui + (1.0 / (t+1.0))

critp = alpha / ui

h = 0

\# Set the counter for false positives to zero j = 0

k2 = 0

s = 0

for y in range(k):

for z in range(x+1):

\# Now create the random numbers per test

a = np.random.normal()

u = st.norm.cdf(a)

pvals[z] = u

p = np.sort(pvals)

\# Now go through each of the values and calculate the number of false positives

j1 = 0

k1 = 0

for i in range(x+1):

\# This is to test for the Bonferroni

if p[i] <= bonthr:

h = h + 1

\# This is for the Benjamini-Hochberg bh1 = ((i+1.0) / (x+1.0)) * alpha

if p[i] <= bh1:

j1 = i + 1

else:

j1 = j1

\# This is for the Narum paper

if p[i] <= critp:

s = s + 1

\# This is for the Benjamini-Yekutieli equation

bh2 = ((i+1.0) / (x+1.0)) * critp

if p[i] <= bh2:

k1 = i + 1

j = j + j1

else:

k1 = k1

k2 = k2 + k1

fwefdr_bonf[x] = ((h * 1.0) / (k * 1.0) * 100.0)

fwefdr_bh[x] = ((j * 1.0) / (k * 1.0) * 100.0)

fwefdr_by[x] = ((k2 * 1.0) / (k * 1.0) * 100.0)

fwefdrnarum[x] = ((s * 1.0) / (k * 1.0) * 100.0)

plt.plot(bonf)

plt.plot(bh)

plt.plot(by)

plt.plot(narum)

plt.show()

